# Bacterial Glycosyltransferase-mediated Cell-surface Chemoenzymatic Glycan Editing: Methods and Applications

**DOI:** 10.1101/322461

**Authors:** Senlian Hong, Yujie Shi, Nicholas C. Wu, Geramie Grande, Lacey Douthit, Ian A. Wilson, Jia Xie, Peng Wu

## Abstract

Chemoenzymatic glycan editing that modifies glycan structures directly on the cell surface has emerged as a complementary tool to metabolic oligosaccharide engineering. In this article, we report the discovery that three bacterial enzymes—*Pasteurella multocida* α2-3-sialyltransferase M144D mutant (Pm2,3ST-M144D), *Photobacterium damsel* α2-6-sialyltransferase (Pd2,6ST) and *Helicobacter mustelae* α1-2-fucosyltransferase (Hm1,2FT)—can serve as highly efficient tools for cell-surface glycan editing. Among these three enzymes, the two sialyltransferases were also found to be tolerant to large substituents introduced to the C-5 position of the cytidine monophosphate *N*-acetylneuraminic acid donor, including biotin and fluorescent dyes. Combining these enzymes with our previously discovered *Helicobacter pylori* α1-3-FT, we developed a live cell-based assay to probe host-cell glycan-mediated influenza A virus (IAV) infection including both wild-type and mutant strains of human H1N1 and H3N2 influenza subtypes. At high SiaNAcα2-6-Gal levels, the ability of a viral strain to induce the host cell death is positively correlated with the SiaNAcα2-6-Gal binding affinity of its haemagglutinin. Surprisingly, the creation of sLe^X^ on the host cell surface *via in situ* α1-3-Fuc editing also exacerbated the killing induced by several wild-type IAV strains as well as a mutant known as HK68-MTA. Structural alignment of HAs from the wild-type HK68 and HK68-MTA revealed the formation of a putative hydrogen bond between Trp222 of HA-HK68-MTA and the C-4 hydroxyl group of the α1-3-linked fucose of sLe^X^. This interaction is likely to be responsible for the better binding affinity of HA-HK68-MTA to sLe^X^ and accordingly the enhanced host-cell killing compared with the wild-type HK68.

## Introduction

Complementary to metabolic oligosaccharide engineering^1^, chemoenzymatic glycan editing has emerged as a valuable tool to modify glycan structures within a cellular environment.^2^ Using a recombinant glycosyltransferase, natural or unnatural monosaccharides can be transferred from activated nucleotide sugars to glycoconjugates on the cell surface with linkage specificity. The ability to install a monosaccharide or its structurally altered analogs linkage specifically to cell-surface glycans provides a facile way for probing its function in a cellular process.

In their pioneering work, Sackstein, Xia, et al., applied chemoenzymatic glycan editing based on human α1-3-fucosyltransferase (FucT) to install α1-3-linked fucose (Fuc) onto the cell surface, thereby creating E-selectin ligand, sialyl Lewis X (sLe^X^, Siaα2-3-Galβ1-4-(Fucα1-3)-Glc*N*Ac), in order to enhance the engraftment and trafficking of human multipotent mesenchymal stromal cells and cord blood cells.^3^ In our previous work, we employed chemoenzymatic glycan editing to tune cell-surface receptor signaling and stem cell proliferation.^2a,4^ Combining this method with bio-orthogonal click chemistry, several labs, including our own, demonstrated that imaging and profiling of specific cellular glycans can be realized.^5^ In a recent, proof-of-concept study, we constructed cell-based glycan arrays using a combination of chemoenzymatic glycan editing, click chemistry, and CHO cells possessing a narrow and relatively homogeneous repertoire of N-linked glycoforms^6^.

To date, glycosyltransferases from both mammalian organisms and bacteria have been used for chemoenzymatic glycan editing. Mammalian glycosyltransferases are type 2 transmembrane proteins.^7^ For cell-surface glycan editing, truncated versions are often used, and include human FucT 6 and 7 expressed in yeast, and ST6Gal1, ST3Gal4, and ST3Gal1 expressed in HEK293 cells.^2a,3a,5b,8^ Bacterial glycosyltransferases, on the other hand, often lack the transmembrane domain and, therefore, are more easily expressed in *Escherichia coli* as soluble proteins. Notably examples include *Helicobacter pylori* α1-3-fucosyl-transferase (Hp1,3FT), the bacterial homologue of the human blood group A antigen glycosyltransferase (BgtA), and the *Campylobacter jejuni* β1-4-N-acetylgalactosaminyl transferase (CgtA).^5c,9^ Surprisingly, we discovered that many bacterial glycosyltransferases that are active for assembly of oligosaccharides in test tubes do not exhibit activities on the cell surface. To expand the enzyme repertoire for chemoenzymatic glycan editing, we performed a screen to identify bacterial glycosyltransferases with relaxed donor specificity that can be used for cell-surface glycan modification.

Here, we report on our finding that three glycosyltransferaes, *Pasteurella multocida* α2-3-sialyltransferase M144D mutant (Pm2,3ST-M144D)^10^, *Photobacterium damsel* α2-6-sialyltransferase (Pd2,6ST)^10,11^ and *Helicobacter mustelae* α1-2-fucosyltransferase (Hm1,2FT)^12^, are able to serve as useful tools for cell-surface chemoenzymatic glycan editing (Figure 1A). Moreover, Pm2,3ST-M144D and Pd2,6ST are tolerant to large substituents introduced to the C-5 position of the CMP-Sia donor, including biotin and fluorescent dyes. We successfully used these two sialyltransferases to survey the expression patterns of their respective glycan acceptors on the surface of live mammalian cells and in tissue specimens. Combining these enzymes with our previously discovered Hp1,3FT, we have developed a live cell based assay to analyze host-cell glycan mediated influenza virus infection.

**Figure 1.**
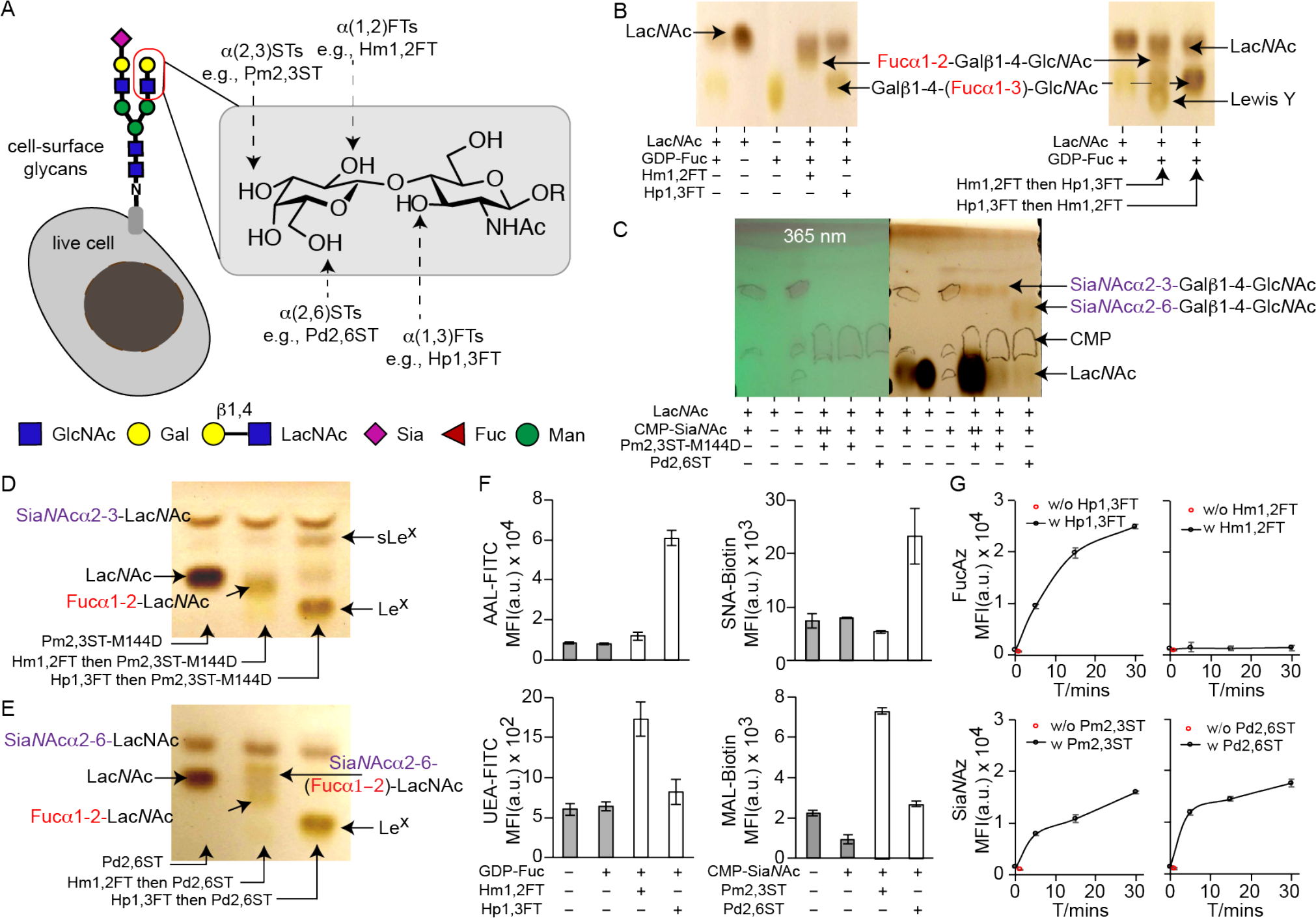
Recombinant bacterial glycosyltransferases (FTs and STs) for live cell-surface glycan editing. (A) An indication of specific positions on mammalian cell-surface Lac*N*Ac(Galβ1-4-Glc*N*Ac)-containing glycans that can potentially be modified by fucosylation (α1-2-or α1-3-linked) and sialylation (α2-3-or α2-6-linked). Recombinant bacterial glycosyltransferases used in this study include Hm1,2FT, Hp1,3FT, Pm2,3ST-M144D and Pd2,6ST. (B) Analysis of *in vitro* fucosylation products by TLC. (C) Analysis of *in vitro* sialylation products by TLC. ++ indicates the final reaction system was further mixed with starting material Lac*N*Ac, and analyzed by TLC. (D and E) Analysis of *in vitro* products generated by a combination of sialylation and fucosylation by TLC. sLe^X^ was formed by combining Hp1,3FT and Pm2,3ST-M144D (D). Sia*N*Ac α2-6-(Fucα1-2)-Lac*N*Ac was formed by combining Hm1,2FT and Pm2,3ST-M144D (E). (F) Analysis of newly formed glycan epitopes on the cell-surface of Lec 2 CHO cells via chemoenzymatic glycan editing. Modified cells were stained by lectins and analyzed by flow cytometry. (G) Evaluation of the substrate tolerance of bacterial glycosyltransferases. Unnatural sugars bearing the azide group were tested, including GDP-FucAz for FTs and CMP-Sia*N*Az for STs. Error bars represent the standard deviation of three biological replicates.

## Results

### Confirmation of the *in vitro* activities of recombinant STs and FTs

Sialic acid (Sia), Fuc and galactose (Gal) are the three most common monosaccharides found on cell-surface periphery glycans. Sia α2-3-or α2-6-linked to terminal Gal, respectively, are exploited by avian and human influenza virus as receptors for host infection. On the other hand, Fuc residues, when attached to terminal Gal in an α1-2-linkage or attached to the Glc*N*Ac of *N*-acetyllactosamine in an α1-3-linkage, form blood group O antigen and Lewis X (Le^X^, Galβ1-4-(Fucα1-3)-Glc*N*Ac), respectively. Unlike Hp1,3FT, which has been used extensively for cell-surface glycan editing, to our knowledge, no other bacterial sialyltransferases (ST) or fucosyltransferases (FT) have been exploited to transfer biophysical probes (e.g. biotin and fluorescent dyes) directly onto cell surfaces for this application.

Chen and coworkers developed the use of two highly efficient enzymes, Pm2,3ST-M144D, and Pd2,6ST, for one-pot chemoenzymatic oligosaccharide synthesis^10,13^. These two enzymes have broad substrate scopes, tolerating functional groups including azide, alkyne, acetyl, *O*-methyl introduced at either the *N*-acyl side chain or the C-9 position. However, it was not known if these enzymes could transfer unnatural Sia analogs functionalized with biotin or fluorescent probes directly onto the cell surface.

We expressed recombinant Hm1,2FT, Pm2,3ST-M144D mutant, and Pd2,6ST as previously described (Figure S1). As a positive control, Hp1,3FT was used. The activities of these enzymes *in vitro* were first verified using the natural donor substrates, CMP-Sia*N*Ac (for STs) and GDP-Fuc (for FTs), and type 2 *N*-acetyllactosamine (Lac*N*Ac) as the acceptor substrate. Thin-layer chromatography (TLC) and liquid chromatography-mass spectrometry (LC/MS) analysis confirmed the formation of Fucα1-2-Galβ1-4-Glc*N*Ac, Siaα2-3-Galβ1-4-Glc*N*Ac, Siaα2-6-Galβ1-4-Glc*N*Ac and Le^X^ in Hm1,2FT, Pm2,3ST-M144D, Pd2,6ST and Hp1,3FT-mediated transformations, respectively (Fig. 1B, C). Consistent with a previous report, when dimeric LacNAc was used as the acceptor substrate, both terminal and internal galactose residues were modified by Pd2,6ST (data not shown).

Subsequently, Fucα1-2-Galβ1-4-Glc*N*Ac was treated with Hp1-3FT or Pd2-6ST to produce Lewis Y or Fucα1-2-(Siaα2-6)-Galβ1-4-Glc*N*Ac, respectively (Figure 1B, E). As reported previously by our lab, sLe^X^ was produced by treating Le^X^ with Pm2,3ST-M144D and CMP-SiaNAc (Figure 1D).

### Evaluation of the activities of recombinant STs and FTs for cell-surface chemoenzymatic glycan editing

We employed the lectin-resistant Chinese hamster ovary (CHO) cell mutant Lec2 that expresses a narrow and relatively homogenous repertoire of glycoforms to examine the feasibility of using the above four enzymes for cell-surface chemoenzymatic glycan editing. The Lec2 cell line has an inactive CMP-Sia Golgi transporter. As a result, it has minimum levels of sialylation, resulting in un-capped Lac*N*Ac and polyLac*N*Ac on cell-surface *N*-glycans. After incubating the cells with each of these enzymes individually along with their corresponding donor substrates, newly formed cell-surface glycan epitopes were probed using fluorescently labeled lectins, including *Ulex Europaeus* agglutinin 1 (UEA 1, specific for α1-2-linked Fuc), *Aleuria Aurantia* lectin (AAL, specific for α1-3-and α1-6-linked Fuc), *Maackia Amurensis* lectin (MAL, specific for α2-3-linked Sia, and *Sambucus Nigra* lectin (SNA, specific for α2-6-linked Sia), respectively. Robust cell-surface lectin staining signals were detected in each set of the experiments (Figure 1F).

Next, we profiled the tolerance of those enzymes for unnatural donor substrates. Azide bearing unnatural donors were tested first, including the fucosylation donor, GDP-FucAz, and the sialylation donor, CMP-SiaNAz. In this experiment, Lec2 cells were incubated with a sialyltransferase (Pm2,3ST-M144D or Pd2,6ST) and CMP-Sia*N*Az, or with a fucosyltransferase (Hp1,3FT or Hm1,2FT) and GDP-FucAz. Following the enzymatic treatment, the modified cells were reacted with an alkynyl biotin *via* the ligand (BTTPS)-assisted copper-catalyzed azide-alkyne [3+2] cycloaddition reaction (CuAAC)^14^, and probed with Alexa Fluor 488-streptavidin conjugate. Flow cytometry analysis revealed that Pm2,3ST-M144D or Pd2,6ST treated Lec2 cells were robustly labeled, and the labeling was time-dependent (Figure 1G). As expected, Hp1,3FT-treated cells also exhibited significant labeling. However, no fluorescent signals were detectable for the Hm1,2FT treated cells, suggesting that this enzyme is unable to accept the azide-functionalized donor. The non-tolerance of unnatural donors by Hm1,2FT was further confirmed by *in vitro* Lac*N*Ac modification. TLC and LC/MS analysis could not detect the formation of the corresponding Fucα1-2-Galβ1-4-Glc*N*Ac derivatives upon incubating Lac*N*Ac with Hm1,2FT in the presence of GDP-FucAl or GDP-FucAz, respectively (Figure S2).

Further evaluation of the donor substrate scope of Pm2,3ST-M144D and Pd2,6ST revealed that besides the *N*-acyl modified CMP-Sia*N*Az, these two enzymes were capable of incorporating other CMP-Sia analogs, including CMP-9AzSia, CMP-SiaNAl and CMP-SiaNPoc, onto cell-surface glycans (Figure S3).

After confirming that Pm2,3ST-M144D and Pd2,6ST could accept unnatural azide-and alkyne-tagged CMP-Sia analogs, we further surveyed if these two enzymes were capable of transferring biotin-or Cy3-functionalized CMP-Sia derivatives directly to the cell surface. To this end, Lec2 cells were incubated with Pm2,3ST-M144D and Pd2,6ST, respectively, in the presence of CMP-Sia-PEG_4_-Biotin or CMP-Sia-Cy3. The biotinylated cells were probed with Alexa Fluor 647-streptavidin conjugate. The cell-surface fluorescence of streptavidin-labeled or Cy3-labeled cells were then quantified by flow cytometry or examined by fluorescent microscopy. We detected strong fluorescent signal in both Pm2,3ST-M144D and Pd2,6ST treated cells. In control experiments, only background fluorescence was observed for cells treated with CMP-Sia*N*Az-Biotin or CMP-Sia*N*Az-Cy3 in the absence of both sialytransferases. As expected Hp1,3FT also incorporated analogs of GDP-FucAz conjugated with Al-PEG_4_-Biotin or Al-Cy3 (Figure 2A-D). To confirm that these signals were produced from glycoprotein labeling, lysates of treated Lec2 cells and Lec8 cells were collected. Anti-biotin Western blot confirmed that biotin was incorporated into glycoproteins of Lec2 cells (MW 55-250 KD), not the mutant CHO Lec8 cells that lack cell-surface Lac*N*Ac (Figure 2F, G). Moreover, PNGase F releasement of *N*-liked glycans essentially abolished all signal of labeled CHO and Lec2 cells, suggesting that Lac*N*Ac residues in N-linked glycans are the primary targets labeled by these enzymes. However, it is also possible that CHO cells express low levels of extended core 1 and core 2 *O*-glycans. Therefore, there are little acceptor substrates to be modified by ST(Pm2,3ST-M144D or Pd2,6ST).

**Figure 2.**
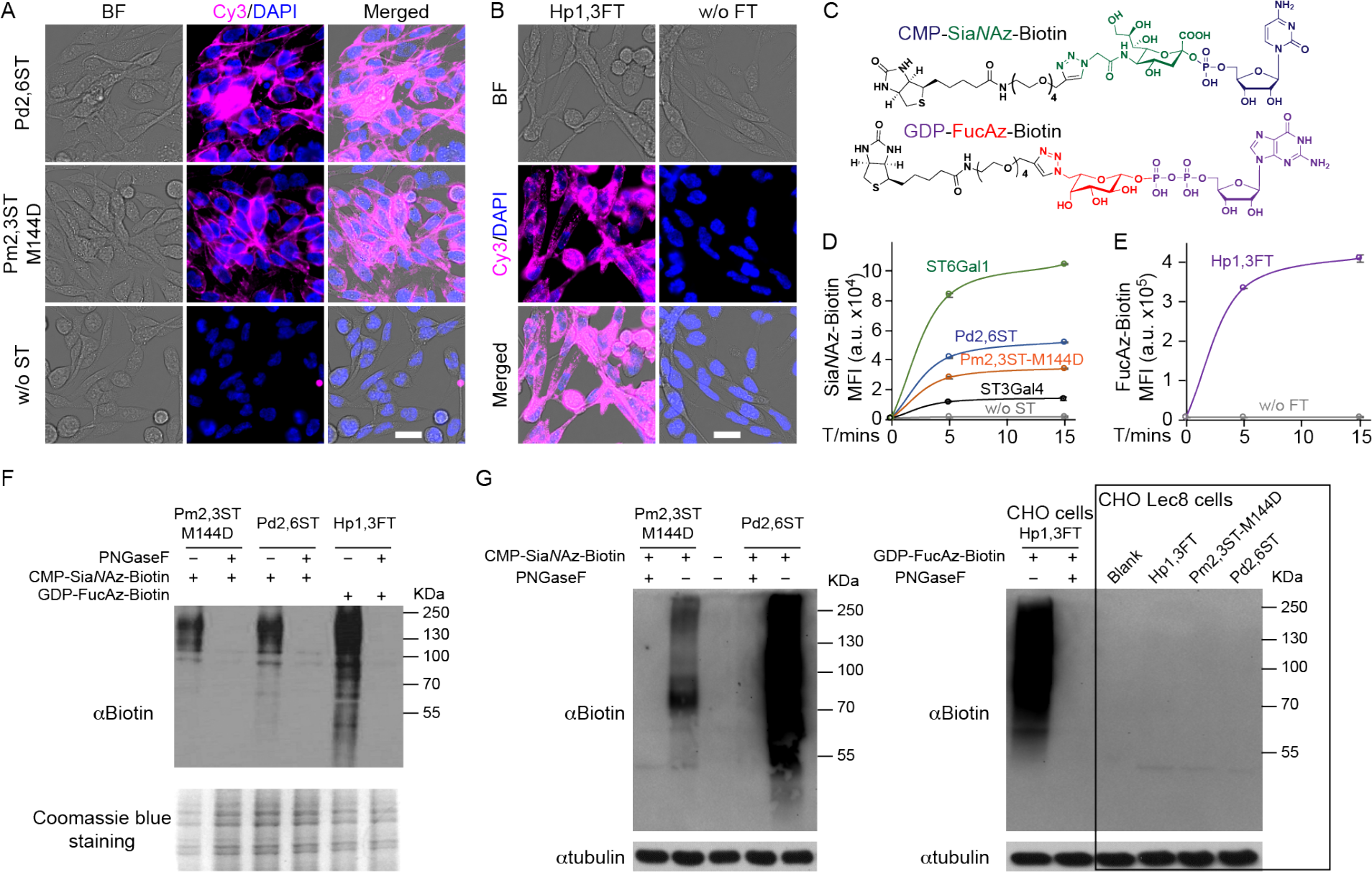
One-step cell-surface glycan labeling enabled by glycosyltransferases (Pm2,3ST-M144D, Pd2,6ST or Hp 1,3ST)-mediated incorporation of unnatural sugars conjugated to a fluorescent dye (Cy3) or an affinity tag (Biotin). (A) Direct STs-catalyzed conjugation of Cy3 (magenta) for imaging of live cell glycans. (B) Hp1,3FT-catalyzed conjugation of Cy3 (magenta) for imaging of live cell glycans. In A and B, cells were visualized by bright field images and DAPI staining (blue). Scale bar, 20 μm. (C) Nucleotide sugars functionalized with biotin tags, CMP-Sia*N*Az-Biotin and GDP-FucAz-Biotin. (D) Time-dependence of activities of recombinant bacterial and human STs for cell-surface glycan labeling with CMP-Sia*N*Az-Biotin. (E) Activity of Hp1,3FT using GDP-FucAz-Biotin to conjugate biotin onto live cell surface glycan directly. In D and E, error bars represent the standard deviation of three biological replicates. (F and G) Enzyme-assisted incorporation of biotin was mainly on N-linked glycans on CHO cells and CHO Lec2 cells, while CHO mutant Lec8 cells that without Lac*N*Ac were not labeled. Protein loading was depicted by Coomassie blue staining or anti-tubulin western blot.

### Labeling of tissue specimens using recombinant bacterial sialyltransferase-based chemoenzymatic glycan editing

Next, we evaluated the feasibility of labeling tissue specimens *via* one-step ST(Pm2,3ST-M144D or Pd2,6ST)-mediated chemoenzymatic glycan editing. Whole embryo frozen sections from C57BL/6 mouse (E16) were used for this evaluation. Embryo sections were incubated with STs (Pm2,3ST-M144D or Pd2,6ST) and CMP-Sia*N*Az-Biotin for 30 mins. The biotinylated samples were then stained with an Alexa Fluor 594-streptavidin conjugate and imaged directly using fluorescent microscopy. Compared to samples without enzyme-treatments, tissue slides treated with STs showed robust fluorescence with distinct labeling patterns (Figure 3, and Figure S4). The outer skin and the salivary gland region exhibited intensive signals afforded by labeling with both enzymes. Interestingly, Pd2,6ST-labeling generated significantly higher signals than Pm2,3ST-M144D-labeling in bone structures, including the sections of leg, rib, spine and skull. In a parallel experiment, tissue sections were digested with PNGase F first to remove *N*-glycans before incubating with STs (Pm2,3ST-M144D or Pd2,6ST) and CMP-Sia*N*Az-Biotin, and probing with an Alexa Fluor 594-streptavidin conjugate. Fluorescence microscopy analysis detected Alexa Fluor 594 associated fluorescence in most organs, strongly suggesting that *O*-glycans are labeled as well (Figure S5).

**Figure 3.**
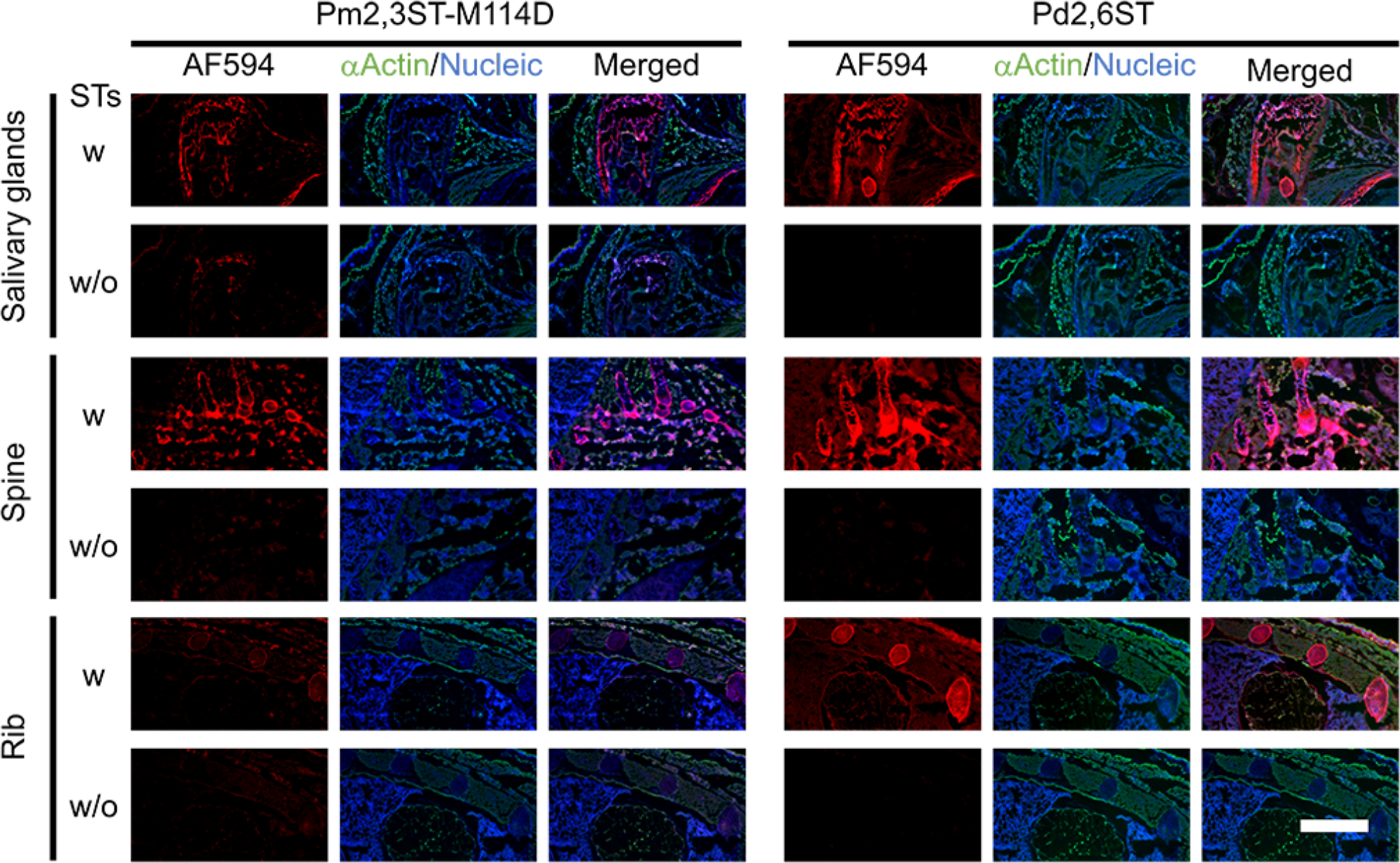
One-step labeling of glycans in tissue specimens of mouse embryos using recombinant bacterial sialyltransferase-based chemoenzymatic glycan editing. The embryonic frozen sections from E16 mouse were incubated with STs (Pm2,3ST-M144D or Pd2,6ST) or without STs, and CMP-Sia*N*Az-Biotin, followed by Alexa Fluor 594-streptavidin conjugate staining. The resulting fluorescence (red) of different parts of the embryo was directly imaged using microscopy, including salivary glands region, lateral sections of spine, and anterior chest. The cells of frozen sections were stained with anti-actin (green) and DAPI (blue, nucleus). Scale bar, 1 mm.

### Construction of a live cell-based glycan array for studying influenza A virus hemagglutinin-glycan interactions

As another application of chemoenzymatic cell-surface glycan editing, we assembled a live cell-based glycan array using the aforementioned enzymes to probe how changes to host cell glycosylation patterns impact influenza A virus (IAV) infection.

The attachment of the hemagglutinin (HA) surface glycoprotein of IAV to the sialylated glycans on the cell surface of host epithelium is the first step in the viral entry cycle.^15^ Glycan microarrays have become heavily employed tools to identify sialylated glycoepitopes that can act as host receptors for IAV and to uncover the Sia binding-preferences of different HAs or whole viruses^16^ It has been found that human IAVs prefer Sia α2-6-linked to Gal (human-type), which is abundantly expressed on epithelial cells of the human airway. By contrast, avian IAVs prefer Sia α2-3-linked to Gal (avian-type) and bind poorly to the human upper airway epithelium.^17^ Despite the rich information gleaned from glycan microarray-based analyses, our understanding of HA-glycan interactions is incomplete without elucidating its physiological relevance. The solid-phase based glycan arrays do not capture the entire potential diversity of glycans present on the cell surface. As revealed by the lectin staining of lung tissues from different donors, cell-surface glycosylation patterns vary from individual to individual, exhibiting fluctuations in α2-3-or α2-6-linked sialylation, α1-3-fucosylation and sLe^X^ expression (Figure 4A, and Figure S6). The variation of glycan expression in a person’s respiratory tract may therefore be responsible for differential susceptibility to influenza infection. We hypothesize that by creating specific glycan epitopes that were previously identified by microarray-based binding assays directly on the live cell surface may serve as a quick way to dissect their specific contributions in a more native environment.

Currently, only two viral subtypes circulate within humans: namely H1N1 and H3N2.^16d^ Using Lec2 cells and a combination of the four enzymes described above, we assembled a small cell-based glycan array displaying various glycan epitopes as shown in Figure 4B. We incubated the HA of influenza A/HongKong/1/1968 (HK68, H3N2) with this array and assessed its binding preference *via* a 3,3’5,5’-tetramethyl benzidine (TMB) assay. As expected, HA of HK68 exhibited strong preference for Sia α2-6-linked to Gal. Surprisingly, it also exhibited significant binding with sLe^X^ created by 1,3FT and 2,3 ST on the cell surface (Figure 4C).

**Figure 4.**
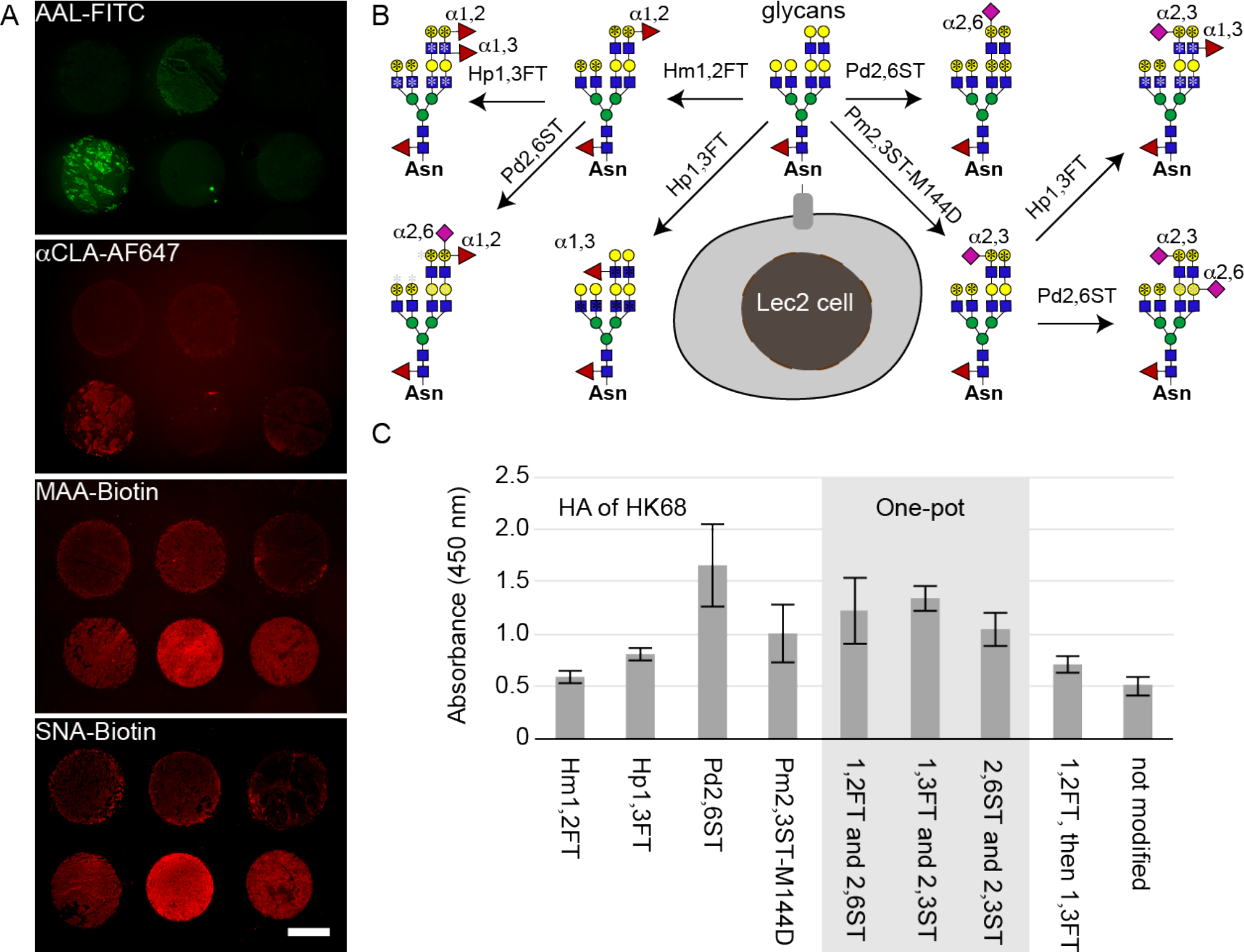
A cell-based glycan array to assess hemagglutinin-glycan interactions directly on the cell surface. (A) Profiling glycoforms of lung tissues obtained from health human donors. Lung tissue slides were stained with FITC-AAL, AF647-anti-CLA, Biotin-MAA or Biotin-SNA conjugates to detect α1-3-fucosylation, sLeX epitopes, α2-3-linked or α2-6-linked sialylation, respectively. (B) Major glycan epitopes presented on Lec2 cell-surface after chemoenzymatic glycan editing. CHO Lec2 cells were treated with glycosyltransferases indicted above and the corresponding nucleotide sugars. *indicate the potential modification site for the first-step glycan editing (black), and the second-step glycan editing (gray). (C) Relative binding affinity of HA from HK68 (H3N2) for glycan-modified Lec2 cells using the specified re-combinant glycosyltransferases. In Figure 4C, the error bars represent the standard deviation of six biological replicates.

### Chemoenzymatic editing of host cell-surface glycans for studying IAV infection

To evaluate if the interaction with sLe^X^ on the host cell surface plays any role in the viral infection, we adopted a live cell-based infection assay. In this assay, we *in situ* edited the glycocalyx of Madin-Darby canine kidney (MDCK) cells, a well-established cell line for studying IAV, using the aforementioned glycosyltransferases. Cultures of the glycocalyx-modified cells, control untreated cells, or cells treated with nucleotide sugars in the absence of any glycosyltransferase, were then infected with serial dilutions of virus in 96-well plates. This assay provides a direct approach to evaluate the impact of Sia and Fuc that are attached to the cell surface with distinct linkages on the susceptibility of host cells to influenza virus infection, enabling correlating glycosylation patterns with host cell killing.

As found in previous studies, both Sia*N*Acα2-6-Gal and Sia*N*Acα2-3-Gal are present on the surface of MDCK cells.^18^ However, the expression level of Sia*N*Acα2-6-Gal is low.^19^ Using Hp1,3 FT-mediated *in situ* Fuc editing, sLe^X^ epitopes can be readily created on the cell surface of MDCK cells, which was confirmed by anti-CLA staining (Figure 5B). Interestingly, all terminal Lac*N*Ac residues on the surface of this cell line have already been capped by Sia. Therefore, no new Sia could be introduced *via* Pm2,3ST-M114D-mediated *in situ* Sia editing (Figure 5A and Figure S7A). This was further confirmed by *in situ* Fuc editing followed by anti-stage-specific embryonic antigen-1 (anti-SSEA-1 or anti-Le^X^) staining, which yielded only background signal (data not shown). However, Gal residues in the internal Lac*N*Ac repeats are still amenable for modification by Pd2,6ST-mediated *in situ* Sia editing to create additional Sia*N*Acα2-6-Gal epitopes (Figure 5A and Figure S7B).

**Figure 5.**
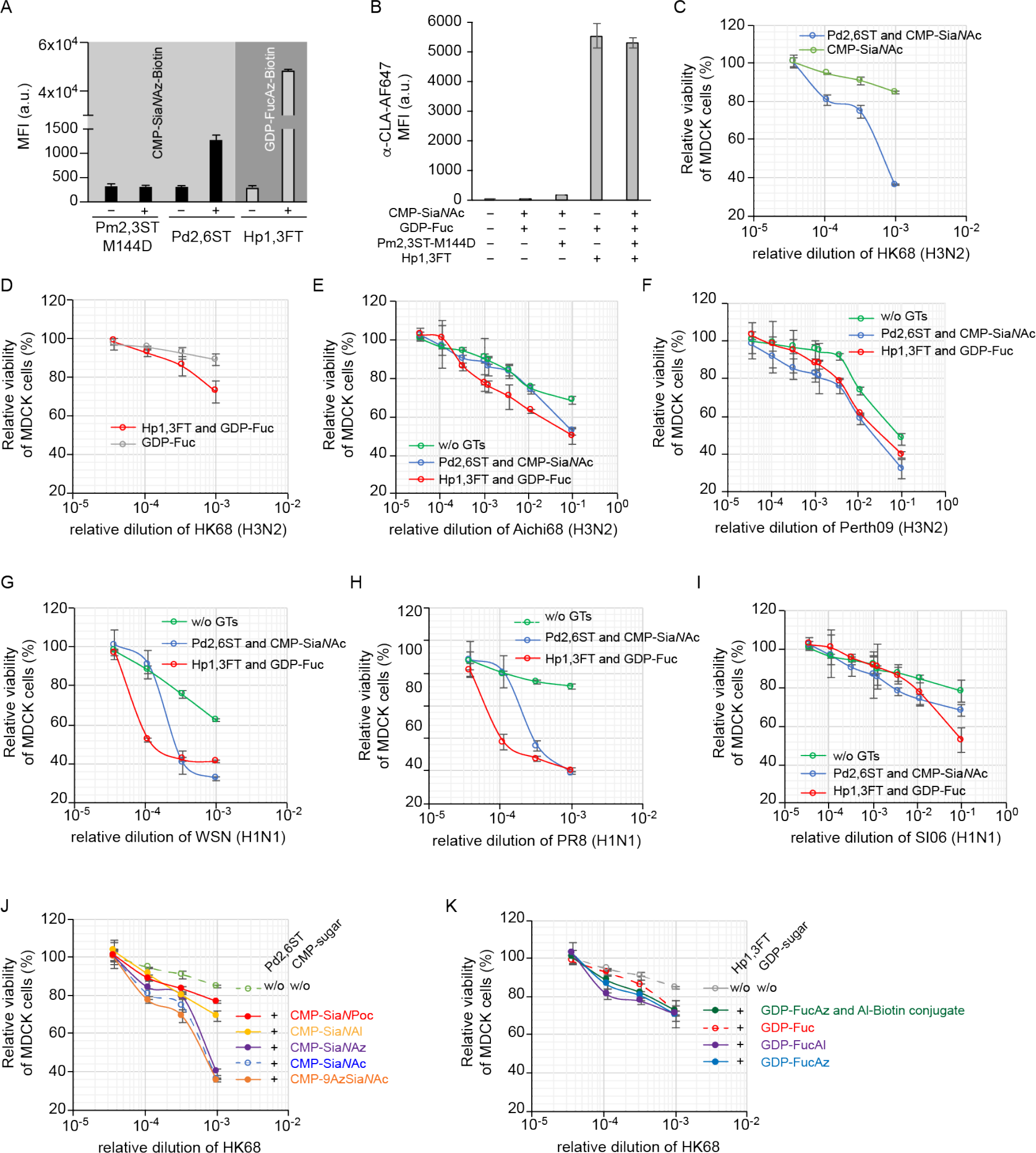
Profiling IAV infection using glycocalyx-modified MDCK cells. (A) Editing glycocalyx of MDCK cells using Pm2,3ST-M144D, Pd2,6ST or Hp1,3ST and the corresponding donor substrate conjugated with biotin. Biotinylated cells were probed with Alexa Fluor 647-Streptavidin. (B) Editing glycocalyx of MDCK cells using a combination of Pm2,3ST-M144D and Hp1,3ST. Newly generated sLe^X^ on the MDCK cell surface was confirmed by Alexa Fluor 647-anti-CLA conjugate staining. (C) Viability of Sia-edited MDCK cells or control cells upon infection by HK68. (D) Viability of Fuc-edited MDCK cells or control cells infected by HK68. (E-I) Viability of glycan (Sia or Fuc) edited MDCK cells or control cells upon infection by Aichi68 (E), Perth09 (F), WSN (G), PR8 (H) and SI06 viruses. (J, K) Viability of glycan edited MDCK cells or control cells upon infection by HK68, using analogs of CMP-Sia (J) or GDP-Fuc (K). In figure A and B, the error bar presented the standard deviation of three biological replicates. In C-K, the error bars represent the standard deviation of six biological replicates.

Naturally occurring H3N2 strains, HK68, A/Aichi/2/1968 (Aichi68) and A/Perth/16/2009 (Perth09), and an H1N1 strain, A/Solomon Islands/3/2006 (SI06), as well as two laboratory-derived H1N1 strains, A/WSN/33 (WSN) and A Puerto Rico/8/1934 (PR8), were used in this infection assay. We first subjected MDCK cells to Pd2,6ST-mediated *in situ* Sia editing or FT-mediated *in situ* Fuc editing to increase cell-surface Sia*N*Acα2-6-Gal epitopes or create new sLe^X^ epitopes, respectively. Next, we incubated the modified MDCK-cells or cells treated with nucleotide sugars only with serial dilutions of influenza viruses. Two days later, the host cell viability was analyzed using 3-(4,5-dimethylthiazol-2-yl)-5(3-carboxymethonypheno l)-2-(4-sulfophenyl)-2H-tetrazolium (MTS) calorimetric assay.

As expected, increasing cell-surface Sia*N*Acα2-6-Gal epitopes enhanced IAV-dependent cell killing for all influenza virus strains tested, especially at high viral titers. In the control experiment, treating cells with the donor substrate CMP-Sia*N*Ac in the absence of Pd2,6ST only had a minor impact on viral infection (Figure 5C-I). Interestingly, the newly added sLe^X^ epitopes on the host cell surface also augmented influenza-induced cell death. At 10^−1^ viral dilution 50±4%, 39±2%, 53±6% of the sLe^x^ decorated cells remained viable upon incubating with Aichi68, Perth09 or SI06 (H1N1), respectively. By contrast, 68±2%, 48±3%, 78±6% of the unmodified cells were viable upon the treatment. More pronounced effects induced by the sLe^x^ addition were observed upon infection with HK68, WSN or PR8; at 10^−3^ viral dilution, only 73±6%, 40±1%, 41±1% of the infected cells remained viable, respectively. When subjected to the same treatment, 89±3%, 62±1%, 92±2% of the unmodified cells were viable.

MDCK cells modified by unnatural Sia and Fuc analogs were also evaluated in this infection assay using HK68. As shown in Figure 5J, although C-9-and *N*-acetyl-Az modified Sia α2-6-linked to Gal exhibited similar activities as the natural ones to promote the influenza virus infection, α2-6-linked SiaNAl and SiaNPoc installed *via* the same fashion showed reduced activities (Figure 5J and Figure S8). Finally, all Fuc analogs examined were found to share similar functions at 10^−3^ virus titer. However, at 10^−4^ virus titer, the alkyne-bearing fucose analog, FucAl, seemed to enhance host-cell infection by HK68 (Figure 5K).

### Host cell-surface glycan editing assists profile the structural flexibility and potential constraints in HA

H3N2 influenza viruses have circulated in humans since 1968, but antigenic drift of HA continues to be a driving force that enables the virus to escape prior immunity. Since most of the major antigenic sites of the HA overlap with the receptor binding site (RBS), the virus constantly evolves to effectively adapt to host immune responses without compromising its virulence.^20^ The RBS consists of the 130-loop, 150-loop, 190-helix, and 220-loop (Wilson et al., 1981).^21^ While the 130-loop, 150-loop, and 190-helix are relatively conserved among HA subtypes, a higher genetic diversity has been detected in the 220-loop, which reflects also some differences in residues responsible for receptor specificity in the different subtypes (e.g. H1N1 vs. H3N2)^16b,16c^.

To examine if sequence variations within the HA-RBS confers H3N2 influenza viruses any advantages to infect host cells harboring Sia*N*Acα2-6-Gal epitopes or sLe^X^ epitopes, we further assessed the wild-type HK68 virus and three laboratory-derived 220-loop mutants that can potentially escape from preexisting immunity by having weaker binding affinity toward the Sia*N*Acα2-6-Gal receptor.^20b^ HK68-MTA (G225M/L226T/S228A), HK68-LSS (G225L/L226S) and HK68-QAS (G225Q/L226A) share a very similar HA backbone conformation, but their binding affinity for Sia*N*Acα2-6-Gal varied significantly.^20b^ Compared with that of wild-type HK68 (WT HK68), the affinities of these mutant HAs to Sia*N*Acα2-6-Gal decrease following the order of HK68-MTA > HK68-LSS > HK68-QAS.^20b^ In the study that created these mutant strains^20b^, all three mutants were found to have WT-like virus replication fitness in unmodified MDCK cells presumably due to the low concentration of α2-6-linked Sia expressed in this cell line.

To assess viral infection in MDCK cells harboring elevated Sia*N*Acα2-6-Gal epitopes or sLe^X^ epitopes, we performed *in situ* Sia or Fuc editing as previously described. The glycan modified cells were then incubated with WT HK68 or the three mutants. The cell viabilities were analyzed after 48 hours. Consistent with previous observations^20b^, all four strains exhibited similar host cell killing capabilities in unmodified MDCK cells (Figure 6D). By contrast, upon elevating the cell-surface Sia*N*Acα2-6-Gal levels, decreased capability to induce host cell death was observed following the order of HK68-MTA > HK68-LSS > HK68-QAS, which matched their Sia*N*Acα2-6-Gal binding affinities. Interestingly, these same mutants manifested different killing capabilities in host cells harboring sLe^X^ epitopes. Compared with WT HK68, enhanced killing was observed for HK68-MTA, whereas HK68-LSS and HK68-QAS exhibited decreased capability to infect sLe^X^-decorated host cells (Figure 6H and 6I).

**Figure 6.**
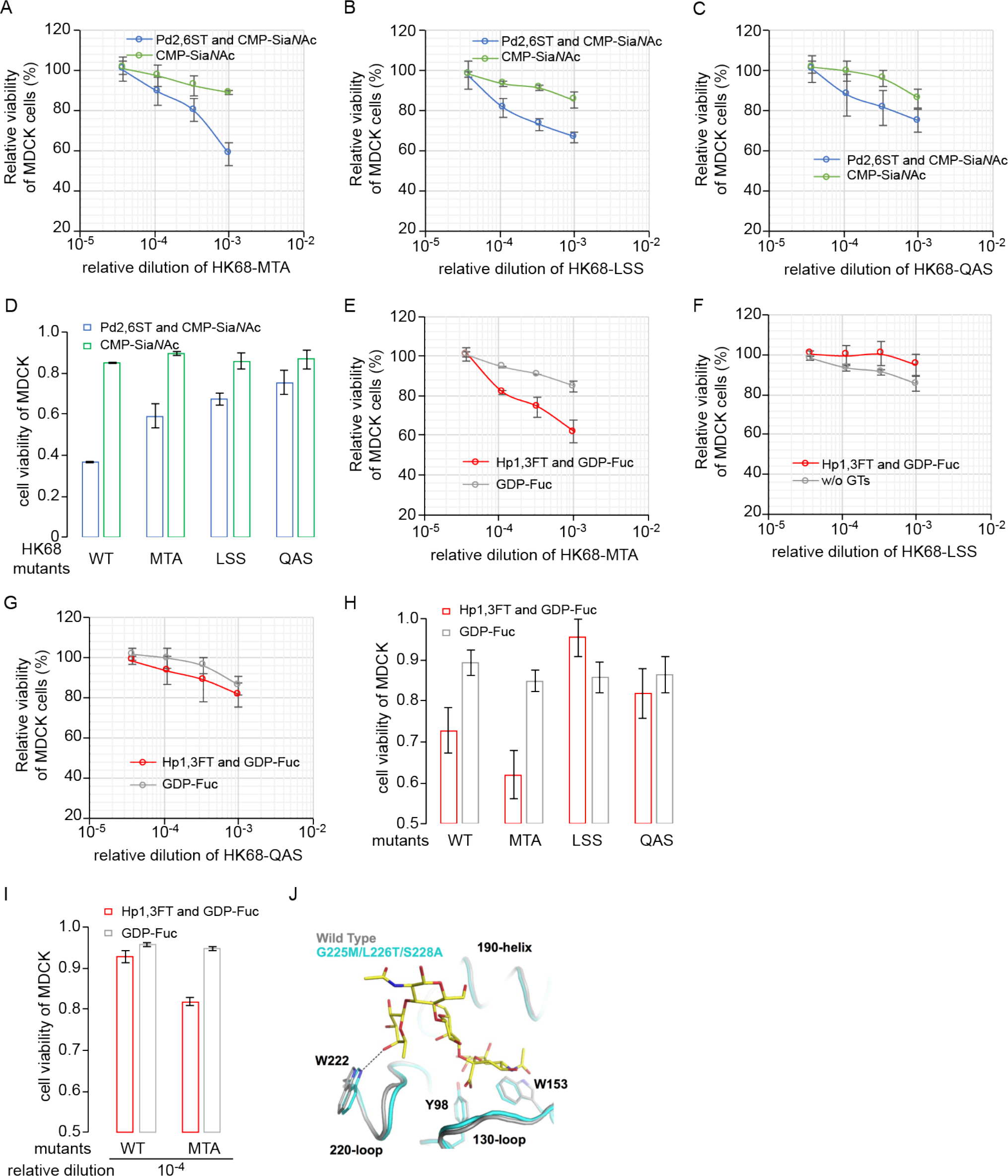
Comparison of the activities of wild-type HK68 and its hemagglutinin-receptor binding site mutants to infect Sia-or Fuc-edited host cells. (A-D) Viability of Sia-edited MDCK cells or control cells upon infection by wild-type HK68 and its HA-RBS mutants, including HK68-MTA (A), HK68-LSS (B) and HK68-QAS (C). (D) Cell viability at 10^−3^ viral dilution. (E-I) Viability of Fuc-edited MDCK cells or control cells upon infection by wild-type HK68 and its HA-RBS mutants. Cell viability at 10^−3^ virus dilution (H), and at 10^−4^ virus dilution (I). (J) Structural alignment of HAs from HK68 and HK68-MTA. A minor shift of 220-loop backbone of HK68-MTA enables formation of a H-bond between C4 hydroxyl of α1-3-linked fucose of sLe^x^ and Nɛ1 of W222 (Figure 6I), which is not observed between the HK68-WT HA and sLe^X^. In Figures 6A-I, the error bars represent the standard deviation of six biological replicates.

Compared with HK68, HK68-MTA was found to possess better preference for sLe^X^ harboring cells, especially at low viral titers (Figure 6H). In order to probe the molecular basis for this observation, apo structures of HK68-WT HA (PDB 4FNK)^22^ and HK68-MTA HA (PDB 5VTX)^35^ were aligned with the crystal structure of A/canine/Colorado/17864/06 (H3 subtype)^23^ HA in complex with sLe^X^ using the RBS (residues 117 to 265 of HA1)^24^ As previously described,^20b^ a minor shift of 220-loop backbone of 0.8 Å was observed in HK68-MTA HA. Our alignment revealed that this shift likely enabled the formation of a H-bond between C4 hydroxyl of Fuc and Nɛ1 of W222 (Figure 6I), which could not be formed between the HK68-WT HA and sLe^X^.

## Discussion

In 1979, Paulson and coworkers demonstrated for the first time that Sia could be directly transferred from CMP-Sia to the cell surface of desialylated erythrocytes using recombinant mammalian sialyltransferases.^25^ Recently, thanks to the creation of the expression vector library encoding all known human glycosyltransferases by Moremen et al, any human glycosyltransferase of interest can now be produced as secreted catalytic domain GFP-fusion proteins in mammalian and insect cell hosts.^26^ Studies by Boons, Steet and coworkers and by our own lab (refs) have demonstrated that several enzymes produced by this system are highly efficient for cell-surface chemoenzymatic glycan editing.^2a,5b,27^ However, this approach is associated with relatively high cost. For cell-surface labeling studies, the GFP tag usually needs to be cleaved before treating cells due to non-specific bindings of GFP to the plasma membrane.

In this study, we discovered that bacterial-derived Pm2,3ST-M144D, Pd2,6ST and Hm1,2FT can be exploited for cell-surface glycan editing. As demonstrated previously and here, these enzymes were easily prepared in multi-milligram quantities in *E. coli* as His-tagged recombinant proteins. Among these three enzymes, Pm2,3ST-M144D, Pd2,6ST were found to tolerate a CMP-Sia donor functionalized with biotin or Cy3, enabling cell-surface acceptor glycans be tagged with these probes for enrichment or visualization. Applying Pm2,3ST-M144D and Pd2,6ST-mediated chemoenzymatic glycan editing to label whole embryo frozen sections from C57BL/6 mice (E16), we found that the salivary gland expressed high levels of acceptor glycans of both enzymes. Sia was first isolated from bovine submaxillary mucin by Blix in 1936. It is not surprising that salivary gland expressed high levels of sialyltransferase acceptors. Interestingly, in the developing bones Pd2,6ST-labeling yielded much higher signals than Pm2,3ST-M144D-labeling. Although Pm2,3ST-M144D can only label the terminal Gal, Pd2,6ST is capable of labeling galactoses of internal Lac*N*Ac units.^11b^ The distinct labeling patents observed here suggest that abundant polyLac*N*Ac glycans are present in bones and in cartilage. This observation is consistent with a previous report that revealed that polyLac*N*Ac were predominantly found in N-glycans of undifferentiated human bone marrow mesenchymal stem cells^28^.

Combined together with our previously discovered *H. pylori* 1,3FT, Pm2,3ST-M144D and Pd2,6ST were used to create a diverse array of sialylated and fucosylated glycan epitopes on the cell surface. By using MDCK cells modified *via* this enzyme-mediated glycan editing to probe the infection of wild-type HK68 and its HA mutants, we confirmed that the ability of an IAV to induce the host cell death is positively correlated to the Sia*N*Acα2-6-Gal binding affinity of the viral HA. Furthermore, this correlation is density dependent—only at high density of cell-surface Sia*N*Acα2-6-Gal, can this correlation be observed. Surprisingly, besides Sia*N*Acα2-6-Gal receptors, several naturally occurring H1N1 and H3N2 strains also recognized sLe^X^ epitopes on the host cells, which facilitated their infection.

HA is the major surface antigen that evolves at an exceptionally high rate. Variation in the HA-RBS through antigenic drift has produced changes in receptor binding that begins to blur the definition of human-type receptor specificity.^20c,29^ Our investigation uncovered that several H3N2 and H1N1 strains, including Aichi68 (H3N2), WSN (H1N1) and PR8 (H1N1), exhibit preference for high sLe^X^-bearing cells over high Sia*N*Acα2-6-Gal-bearing cells especially at low viral titers (Figure 5). At low virus conditions, these strains induced significantly higher levels of cell death in sLe^X^-harboring MDCK cells than in Sia*N*Acα2-6-Gal-harboring counterparts. These observations suggest that such strains may selectively infect human populations with high sLe^X^-expression in their respiratory tracts, such as patients with cystic fibrosis and patients suffering from airway inflammation.^30^ It has been documented that several avian influenza virus strains exhibit strong affinities for sLe^X^-type receptors.^17c,17d,31^ Therefore, it is likely that human populations with high sLe^X^-expression in their respiratory tracts are susceptible to these viruses as well.

Our studies confirmed that binding specificity and strength to HA are not only encoded in the structures of individual glycans, but are also determined by the density of these epitopes on the cell surface. This context-dependent molecular recognition underscores the importance of tools that empower the investigation of glycan functions within a more native environment such as the cell surface. The chemoenzymatic glycan editing technique described here should serve as a valuable tool for accomplishing this goal. Currently, we are applying this technique to explore the impact of changes to cell-surface glycosylation patterns on the infection of other types of human viruses.

